# Evaluation of EV Storage Buffer for Efficient Preservation of Engineered Extracellular Vesicles

**DOI:** 10.1101/2022.12.06.519345

**Authors:** Yuki Kawai-Harada, Hanine El Itawi, Hiroaki Komuro, Masako Harada

**Affiliations:** Institute for Quantitative Health Science and Engineering (IQ), Michigan State University; Department of Biomedical Engineering, Michigan State University

**Keywords:** Extracellular vesicles, storage buffer, preservation, Engineered Extracellular vesicles

## Abstract

Extracellular vesicles (EVs), detectable in all bodily fluids, mediate intercellular communication by transporting molecules between cells. The capacity of EVs to transport molecules between distant organs has drawn interest for clinical applications in diagnostics and therapeutics. Although EVs hold potential for nucleic acid-based and other molecular therapeutics, the lack of standardized technologies, including isolation, characterization, and storage, leaves many challenges for clinical applications, potentially resulting in misinterpretation of crucial findings. Previously, several groups demonstrated the problems of commonly used storage methods that distort EV integrity. This work aims to evaluate the process to optimize the storage conditions of EVs and then characterize them according to the experimental conditions and the models used previously. Our study reports a highly efficient EV storage condition, focusing on EVs’ capacity to protect their molecular cargo from biological, chemical, and mechanical damage. Compared with commonly used EV storage conditions, our EV storage buffer leads to less size and particle number variation at both 4°C and -80 °C, enhancing the ability to protect EVs while maintaining targeting functionality.

## Introduction

First discovered in the 1960s^1^, extracellular vesicles (EVs) are a heterogeneous population of lipid-bound nanoparticles secreted by all cell types in all species.^2^ The role and secretion mechanism were obscure until two separate groups simultaneously reported the selective release of small vesicles containing transferrin receptors that was internalized to reticulocytes in 1983.^3,4^ They revealed the formation of exosomes as intraluminal vesicles of multivesicular endosomes that are secreted extracellularly upon fusion of the endosome and plasma membrane.^3^ In the last decades, EV (including exosome) function in cell-cell communication became clearly evident and thus has received increasing attention. Numerous groups reported the function of EVs in a broad range of physiological and pathophysiological processes, including immune responses^5^, neurodegenerative diseases^6^, inflammatory diseases^7^, cardiovascular diseases^8^, cancers^9^, and infectious diseases^10^, which were mediated by transferring functional biomolecules, such as protein, nucleic acids, and metabolites. EVs are an attractive source for diagnostic biomarkers because methods of non-invasive body fluids sampling are mostly well-established. In addition, the properties of EVs to protect and transport bioactive molecules are ideal as a delivery vehicle for molecular drugs for the current molecular medicine needing carriers that do not elicit immunogenic responses. However, the progress toward clinical use of EVs is slow due to the shortfall of standardized methods, which often result in experiment-to-experiment variabilities and contradictory findings on their biology and functions, impeding the progress of EV research, not limited but including isolation, characterization, cargo loading and controlled and mass production of EVs.^11^ Therefore, the International Society for Extracellular Vesicles (ISEV), continues to update the essential Information for Studies of Extracellular Vesicles (MISEV) guidelines.^12-14^

An isolation method is one of the critical processes in EV research, and includes differential ultracentrifugation, gradient centrifugation, filtration, affinity-immunoprecipitation, and microfluidic isolation. Each method allows the enrichment of slightly different EV populations but is associated with a different grade of impurity, which also depends on the EV source (culture media or bodily fluids).^11,15^ Another area urgently needed in EV research is the physical and chemical properties of storage conditions that influence EV quality for both analysis and downstream applications.^16,17^ The most commonly used freeze storage in phosphate buffer saline (PBS) causes damage to EVs during storage leading to the loss of some functions.^16,18^ Several groups explore various conditions and the use of additives to improve the conditions, such as the use of Tween^18^, BSA^19^, trehalose^20^, and DMSO^19^, at different temperatures. A recent report by Görgens *et al*. showed that adding the combination of Human Serum Albumin (HSA) and trehalose to PBS-HEPES buffer drastically improved EV quality following short- and long-term storage compared to PBS alone.^21^ Interestingly, a widely-used buffer additive, serum albumin is the most abundant protein component of the blood of all vertebrates, which has evolved by diversifying the physicochemical, genetic, and physiological biochemical properties, thus variable in different species.^22^ While HSA possesses extensively overlapping physicochemical properties, it differs from BSA, for example, in adsorption, crystallization mechanisms, and binding affinity to hydrophilic surfaces.^23^ Thus, a careful evaluation of each serum is necessary as these properties influence interactions.^24^

In this study, we evaluated the EV storage buffer (EBV), consisting of trehalose, BSA supplemented PBS-HEPES buffer, compared to the widely used storage methods (PBS or media storage) for the capacity of engineered EVs, focusing on cargo protection, which is one of the critical features in drug/gene delivery system. Our study complements the previous report by Görgens et al. that uses HSA instead of BSA and validates the preservation method using the more readily available BSA, but includes additional parameters.^25^ The use of protected EV cargo, rather than the total EVs as the primary independent assessment, can provide a more precise degree of the transportation capacity of EVs within mammalian systems.

## Materials and Methods

### Cell Culture and Treatment

293T (Human Embryonic Kidney cell line), were obtained from American Type Culture Collection (ATCC) and tested for mycoplasma. The cells were cultured in high-glucose DMEM (Gibco) supplemented with 100U/mL penicillin, 100μg/mL streptomycin and 10% (v/v) fetal bovine serum (FBS, Gibco). Engineered EVs were generated by transfecting EV-display plasmids (pcS-mCherry-C1C2 for storage test, pcS-E626-C1C2/pcDNA-gLuc-C1C2 for in vitro testing, and pcS-p88-C1C2/pcDNA-gLuc-C1C2 for in vivo testing.) into 293T cells using home-made PEI as described previously.^26,27^ Following 24 h incubation, cells were washed twice with PBS to remove residual PEI-DNA complex and EVs derived from FBS, and the culture media was replaced with DMEM supplemented with Insulin-Transferrin-Selenium (ITS) Growth Supplement (Corning), 100U/mL penicillin, 100μg/mL streptomycin for another 24 h incubation for EV production. The clones are available either from the addgene (https://www.addgene.org/Masako_Harada/) or the corresponding author upon request.

### EV Isolation

The cells were grown in DMEM media supplemented with ITS and Pen-Strep for 24 h, and the media from the plates was collected. For each batch, EVs were purified from 20 mL of conditioned media by differential centrifugation. The media was centrifuged at 600g for 30 min to remove the cell and cell debris. In order to remove the contaminating apoptotic bodies, the media was centrifuged at 2000g for 30 min. The supernatant was then ultracentrifuged in PET Thin-Walled ultracentrifuge tubes (Thermo Scientific 75000471) at 100,000g with a Sorvall WX+ Ultracentrifuge equipped with an AH-629 rotor (k factor = 242.0) for 90 min at 4 °C to pellet the EVs. The pellet containing EVs was resuspended in 100 μL PBS or the EV buffer. Filter sterilized EV buffer consists of 0.2% Bovine Serum Albumin (Fisher BioReagents™), 25mM D-(+)-Trehalose dihydrate (TCI America), 25mM HEPES pH7.0 (Sigma) in PBS pH7.4 (Gibco).

### Nanoparticle Tracking Analysis (NTA)

The particle size and concentration were measured using a ZetaView® (Particle Metrix) Nanoparticle Tracking Analyzer following the manufacturer’s instruction. The following parameters were used for measurement (Post Acquisition parameters [Min Brightness: 22, Max area: 800, Min area: 10, Tracelength: 12, nm/Class: 30, Classes/Decade: 64]) (Camera control [Sensitivity: 85, Shutter: 250, Frame Rate: 30]). EVs were diluted in PBS between 20- and 200-fold to obtain a concentration within the recommended measurement range (0.5×10^5^ to 10^10^ cm-^3^).

### Immuno-Transmission Electron Microscopy (Immuno-TEM)

EV particle number were determined by NTA for each storage condition. Carbon film coated 200 mesh copper EM grids were soaked in 50 μL EVs (1×10^7^ mCherry EVs in PBS or EV storage buffer) for 30 min for the adsorption of EVs on the grid. EVs on the grids were fixed by treating with 50 μL of 2% Paraformaldehyde (PFA) for 5 min and then rinsed thrice with 100 μL PBS. To quench free aldehyde groups, the grids were treated with 50 μL of 0.05 M glycine for 10 min. The surface of the grids was blocked with a drop of blocking buffer (PBS containing 1% BSA) for 30 min. After blocking, the grids were incubated with 50-100 μL anti-HA (Sigma-Aldrich H3663) or anti-CD63 (Thermo Fisher 10628D) antibody (1:100 in PBS containing 0.1% BSA) for 1 h. The grids were washed five times with 50 μL PBS containing 0.1% BSA for 10 min each. For secondary antibody treatment, the grids were incubated in a drop of Goat-anti-Mouse IgG coupled with 10 nm gold nanoparticles (Electron Microscopy Sciences, 25512) diluted at 1:100 in PBS containing 0.1% BSA for 1 h. The grids were washed five times with 50 μL PBS containing 0.1% BSA for 10 min each and then with two separate drops of (50 μL) distilled water. EVs were negatively stained with 2% uranyl acetate and then rinsed with PBS. The grids were then air dried for 24 h and images were captured by transmission electron microscope (JEOL 1400) at 80 kV.

### DNase I Treatment and Plasmid DNA recovery from EVs

The 2 μL of EVs were incubated at room temperature for 15 min with 1 U of DNase I (Zymo Research) and 1x DNA Digestion Buffer. The plasmid DNA was isolated from the EVs using Qiamp Miniprep kits and quantified by qPCR. The plasmid DNA was also extracted from Non-treated EVs as a negative control.

### Quantitative Real-time Polymerase Chain Reaction (qPCR)

qPCR was performed using DreamTaq DNA polymerase (Fisher BioReagents). Each reaction contains 200 μM dNTP, 500 nM each of forward (5’-CTAGAGTAAGTAGTTCGCCAGTTAAT-3’)/reverse primer (5’-GCTGAATGAAGCCATACCAAAC-3’), 200 nM probe (5’-ATTGCTACAGGCATCGTGGTGTCA-3’), 0.5 U DreamTaq DNA polymerase, 1x DreamTaq buffer and 1 μL sample DNA in a total reaction volume of 10 μL using CFX96 Touch Real-Time PCR Detection System (BIO-RAD). The PCR amplification cycle was as follows: 95°C for 2 min; 40 cycles of 95°C for 20 seconds, 65°C for 30 seconds. The plasmid DNA (pDNA) copy number was determined by absolute quantification using qPCR to calculate the copy number of EV-encapsulated pDNA per vesicle based on NTA.

### In vitro Bioluminescence Binding Assay

Engineered EVs were generated by co-transfecting EV-display gLuc plasmid (pcDNA-gLuc-C1C2) and EV-display-E626 plasmid (pcS-E626-C1C2) into 293T cells for bioluminescence binding assay as previously described.^27^ Particle numbers for each condition were measured right before the binding assay following storage. Briefly, A431 cells were seeded at 2.0×10^4^ cells/96-well TC plates 24 h prior to EV treatment. The cells were treated with 2.0 × 10^7^ particles of E626-gLuc-EVs in 100 μL media for 0, 10, 30, and 60 min at 37 °C. Following the three PBS washes to remove unbound EVs, 1 μg/mL coelenterazine-H (CTZ; Regis Technologies, Morton Grove, IL, USA) in 95 μL of DPBS was added to the wells and imaged by an in vivo imaging system (IVIS; Spectrum Perkin Elmer, Aaltham, MA, USA). Total photon flux (photons/s) was quantified using Living Image 4.7.2 software (IVIS, PerkinElmer). Values are presented as the means ± SD (n = 4).

### Animals and Ex vivo Bioluminescence Biodistribution Assay

Adult female Balb/cJ mice weighting 18-20 g (7-8 weeks old) were used for animal experiments. Animals were purchased from Jackson Laboratories and housed in the University Laboratory Animal Resources Facility at Michigan State University. All the experimental procedures for the animal study were performed with the approval of the Institutional Animal Care and Use Committee of Michigan State University. Anesthetized mice received intravenous injection of 1.0×10^10^ p88-gLuc EVs stored for 24 h-post isolation either in DPBS or EV buffer. Following 60 min circulation, the mice were sacrificed, and the following visceral organs were dissected and placed on a transparent sheet: heart, lungs, liver, kidneys, spleen, and pancreas. Ex vivo bioluminescence images were taken following the application of 200 μL CTZ (10 μg) to each organ, and the total flux (photons/s) of images was quantified using Living Image 4.5 software (IVIS, PerkinElmer). Values are presented as the means ± SD (n = 3).

### Data analysis and statistics

One-way ANOVA and t-test were performed to assess the effect of time and storage conditions. Further details are provided in respective figure legends. Graphs were assembled using GraphPadPrism9 (GraphPad Software).

## Results

### Experimental design and generation of engineered EVs (eEVs) for storage test

The experiments were designed to evaluate the short-term storage condition for engineered EVs compared to PBS and culture media as control. Engineered EVs with modified surfaces that package plasmid DNA were subjected to storage using various buffers and temperature conditions for up to 7 days (Table 1). The formulation and condition of the EV storage buffer were carefully selected based on the addition of trehalose^20^ and BSA-HEPES^18^ reported by Bosch *et al*. and van de Wakker *et al*., respectively, which overlaps with the PBS-HAT buffer described by Görgens *et al*. (Note that the sources of albumin are different.)

**Table 1.**
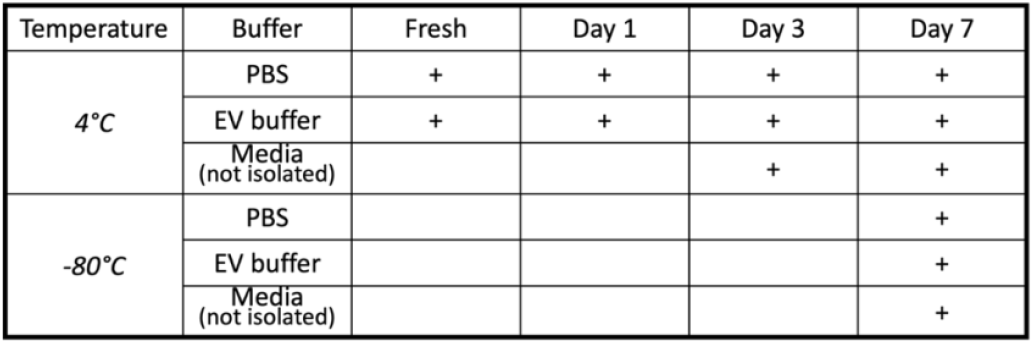
The evaluated timepoint, storage temperature, and conditions.

Engineered EVs were generated as previously described using genetic modification to the HEK293T cells and cultured media isolation by differential ultracentrifugation (Fig 1)^26,27^ following the pellet resuspension in each buffer. Fresh samples were subject to each assay within 2 hours following the isolation. These EVs were previously characterized and validated for EV surface display and pDNA packaging using NTA, western blotting, immuno-Transmission Electron Microscopy (Immuno-TEM), and qPCR.^26,27^

**Figure 1.**
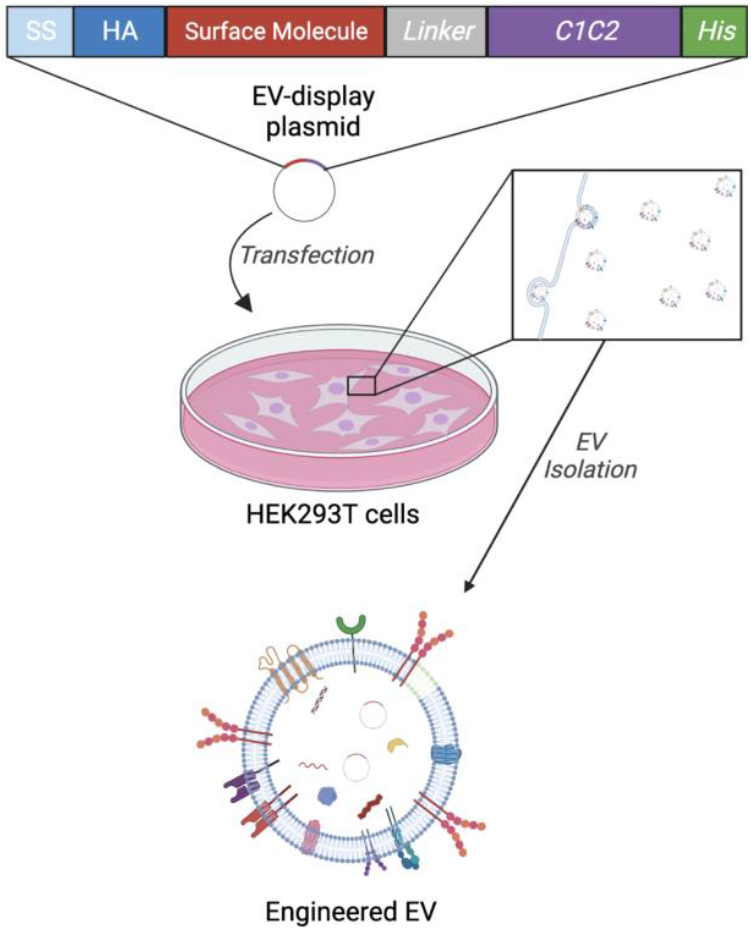
Schematic diagram of the eEV preparation. The EV generation from HEK293T cells by transfecting the EV-display pDNA and secretion of engineered EVs into the cultured media, resulting in the generation of engineered EVs. (Created with BioRender.com)

### EV storage condition influence recovered particle numbers and sizes

First, we directly observed the size and morphology of eEVs using Immuno-TEM, which showed heterogeneity in size and morphology for each condition (Fig S1). Then, we measured particle concentrations and sizes for each condition using Nanoparticle Tracking Analysis (NTA). As shown in figures 2A, B, and C, PBS-stored eEVs had significant particle reductions over time, while EV buffer-stored eEVs did not show a statistically significant difference after the 7-day storage period. The particle numbers between fresh samples stored in PBS and the EV buffer showed no statistical significance. Storage of EVs without isolation (cultured media) consistently led to the recovery of fewer particles regardless of the storage temperature, suggesting that this storage method caused a more significant loss of particles than isolated EV storage (Fig 2D).

**Figure 2.**
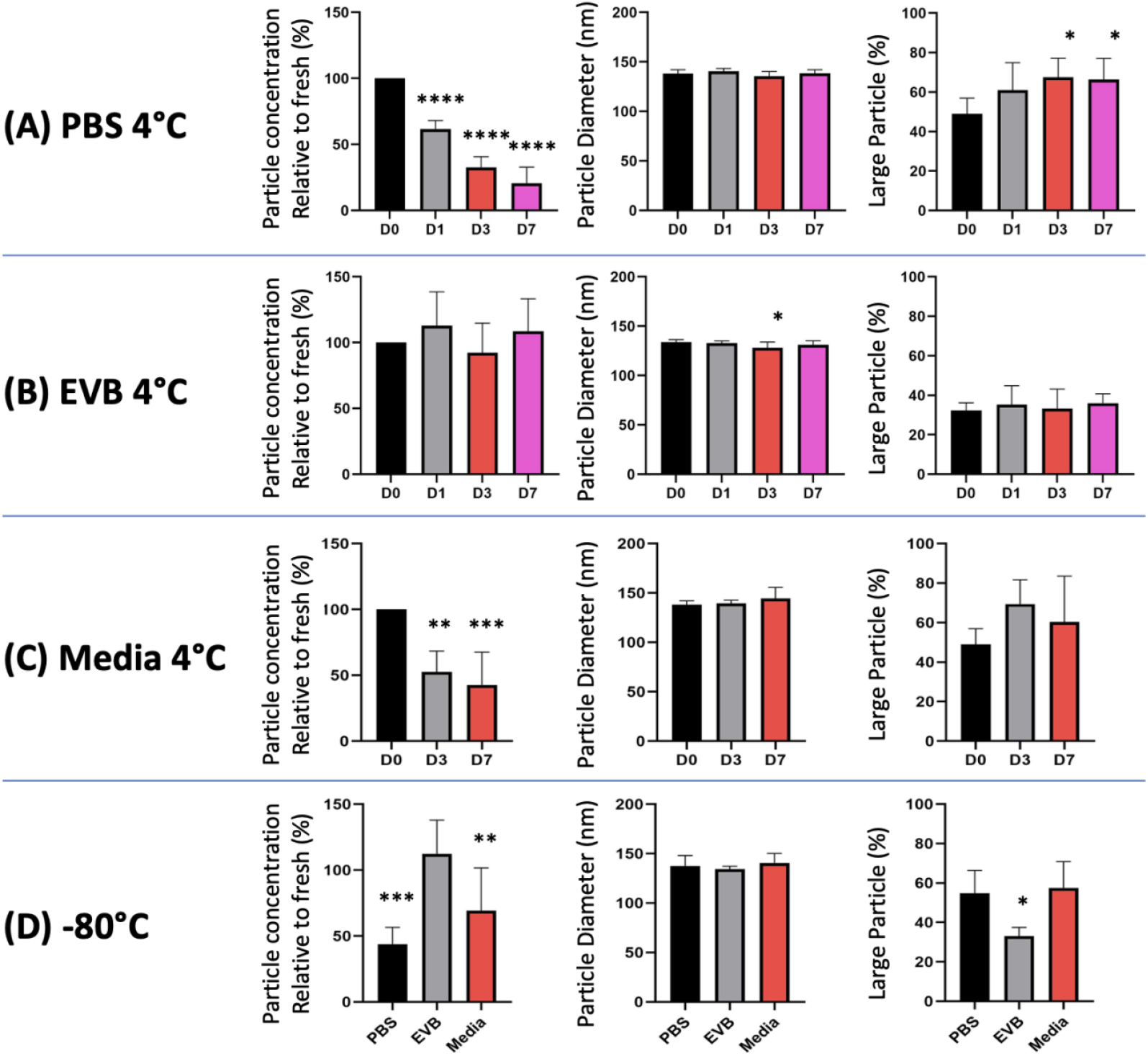
Effects on eEV concentration and sizes after eEV storage in PBS and the EV buffer. eEV were isolated from HEK293T conditioned medium by differential ultracentrifugation, re-suspended in (A) PBS, the (B) EV buffer, or (C) Media without isolation and stored at both 4°C or (D) -80 °C for indicated duration. Effects of storage on EV concentration, size, and volume distribution are shown. Particle number was standardized on yield from 1 mL culture media. The data of PBS Day 0 was used as the data of media day 0. (n=7) One-way ANOVA was used to evaluate the effect of the time course in the group, unpaired t-test was used to compare different storage conditions. In all figures, significance is expressed as follows: *P≤0.05, **P≤0.01, ***P≤0.001, ****P≤0001, if not otherwise specified.

To examine the influence of EV storage on eEV sizes, we compared the diameter of eEVs and found no significant differences in the median peak across samples (Fig 2A, B, and C). However, there was a general trend toward the increased volume of larger particles in PBS or Media storage compared to the EV buffer, indicated in the Volume/mm3E-6 graph (Fig S2), which was also observed visually in the video capture (data not shown). Therefore, we further analyzed these data by selecting the particles larger than 200 nm and plotting the % volume for each condition, which exhibited a significant increase in large particle volume within the PBS-stored eEVs (Fig 2A). Notably, the difference in the large EV volumes between samples stored in PBS and EV buffer at -80°C (Fig 2D) suggested that the EVs in preparation started fusing or aggregating immediately after re-suspension in PBS.

### The EV storage conditions influence the integrity of eEVs and packaged DNA contents

One of the crucial aspects of eEV storage is its influence on the protective properties of eEVs. To address this question, following the comparison of the physical properties of EVs, we isolated pDNAs from eEVs using qPCR before and after the DNase I treatment to examine the change in eEVs’ protective properties. Notably, freezing media (unisolated eEVs) or eEVs stored in PBS reduced EV protection significantly compared to eEVs stored in the EV buffer. In contrast, there was no significant difference in pDNA protection at 4 °C in both buffers (Fig 3A). We further analyzed pDNA copy numbers per EVs based on the qPCR and NTA data and found no substantial differences among storage conditions (Fig 3A and S2). However, when we calculated based on the original media volume by converting to the uL of EV suspension, pDNA copy numbers were reduced both in PBS and -80 °C (Fig 3B). These data implicate total loss of eEVs in PBS and destruction of eEVs by -80 °C storage. Most importantly, the EV buffer retained higher EV numbers and pDNA copies following either 4°C or 80 °C over the 7-day storage period.

**Figure 3.**
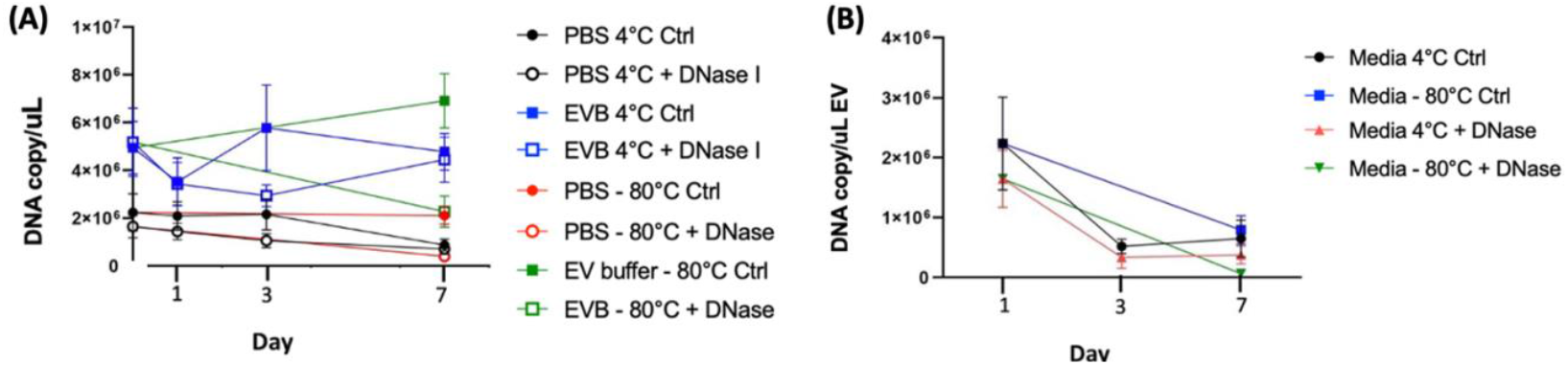
Effects of storage on pDNA protected in eEVs. eEV were isolated from HEK293T conditioned medium by differential ultracentrifugation, re-suspended in PBS or the EV buffer and stored at both 4°C or -80 °C for indicated duration. Effects of storage on **(A)** pDNA protection rate, **(B)** pDNA copy number per single EV, and **(C)** pDNA copy numbers per 1 μL of EVs are shown. The data of PBS Day 0 was used as the data of media day 0. (n=4) One-way ANOVA was used to evaluate the effect of the time course in the group, unpaired t-test was used to compare different storage conditions. In all figures, significance is expressed as follows: *P≤0.05, **P≤0.01, ***P≤0.001, ****P≤0001, if not otherwise specified.

### The effect on the eEV functionality by EV buffer storage

Another critical aspect of eEV storage is whether to influence the function as a delivery vehicle, especially its targeting capacity. To test the storage effect, we assessed the binding capacity of eEVs by bioluminescence imaging (BLI) using a previously established system that demonstrated EGFR-targeting EVs.^27^ Briefly, EGFR-overexpressing cells were treated with eEVs co-labeled with Gaussia luciferase (gLuc) and an EGFR-targeting monobody, washed, substrate added, and binding measured by BLI. As shown in figure 4, the bioluminescence from bound eEVs was significantly improved in EV buffer compared to PBS. However, the bioluminescence in the EV buffer was significantly higher (Fig S3), possibly due to the influence of the buffer formulation, suggesting the binding was not enhanced but at least not interfered with. It is supported by the in vivo data in the following.

**Figure 4.**
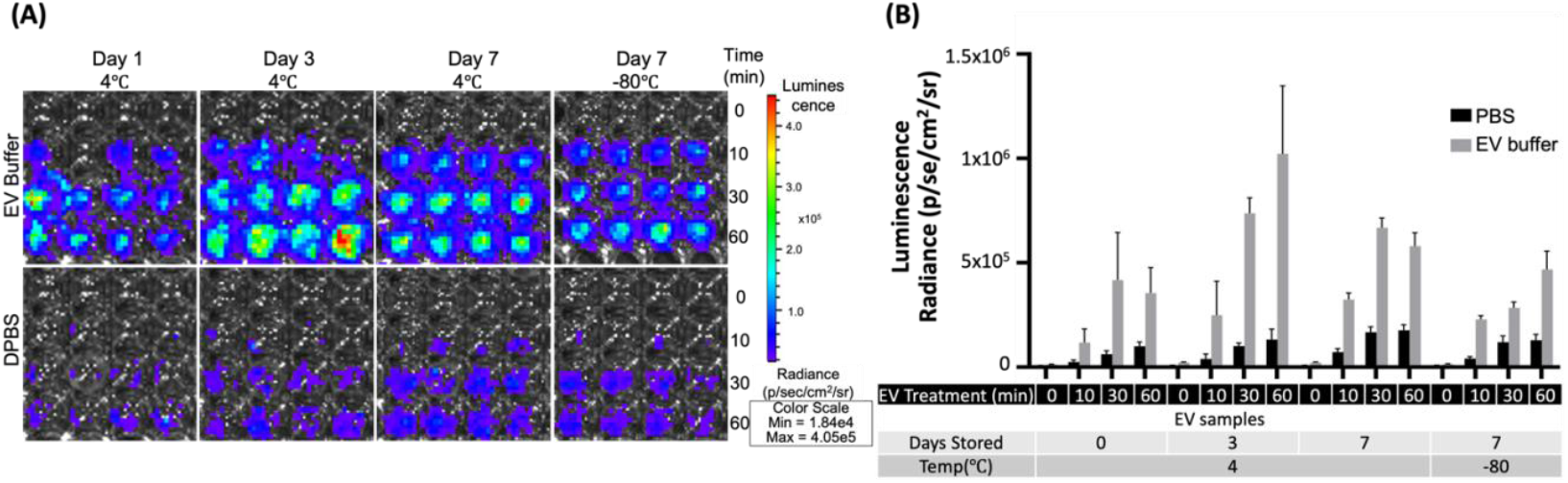
Effects on eEV binding capacity after eEV storage in PBS and the EV buffer. A431 cells (EGFR-positive cells) treated with targeting (E626) EVs. **(A)** Representative image of eEV (gLuc) binding to A431 cells after eEV storage in PBS and the EV buffer for the indicated time. **(B)** The total photon flux (p/s) from EVs bound to the cells by IVIS. The value represents the means ± SD (n = 4) in the graph.

Further, we used a pancreas-targeted EV display system to assess differences in vivo function after storing 24 h at 4 °C in PBS or EV buffer.^26^ Briefly, eEVs co-labeled with gaussia luciferase (gLuc) and a pancreas-targeting peptide (p88) were injected into a mouse via tail vein, circulated for an hour, sacrificed, and organs removed for BLI measurement. As shown in Figures 5A and B, there were no significant differences in targeting capacity or signal intensity, consistent with our previous findings,^26^ indicating the storage buffer does not influence the targeting capacity of eEVs *in vivo*.

**Figure 5.**
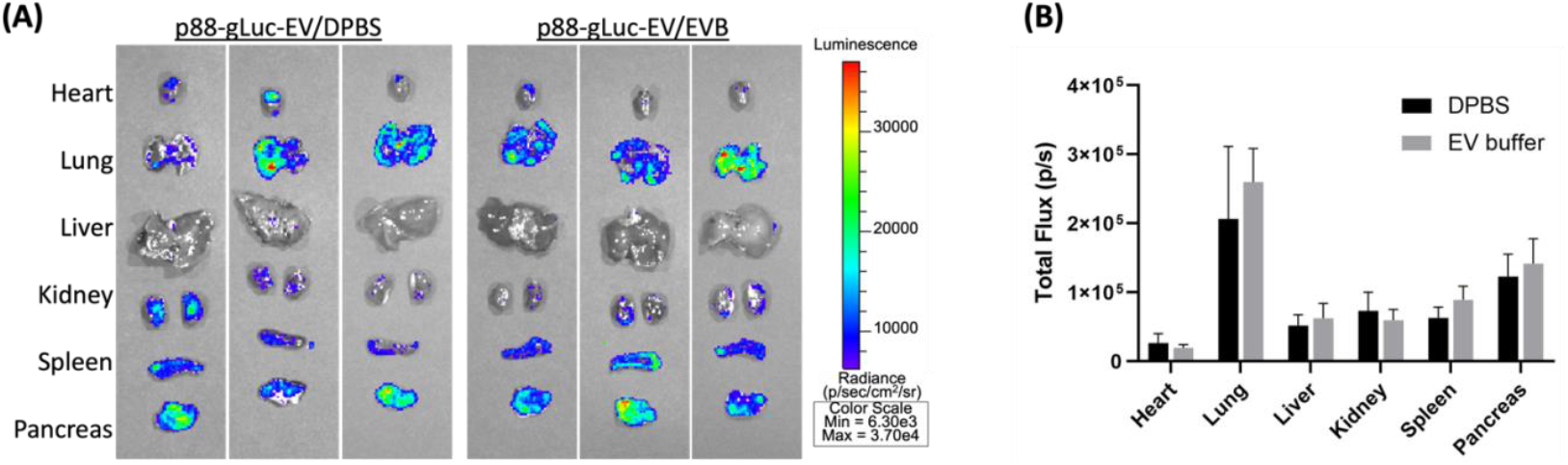
Comparison of Biodistribution by peptide-displayed eEVs stored in PBS and the EV buffer (A) Representative images of the organs from the Balb/cJ mice received intravenous injections of p88-gLuc-EVs stored in PBS and the EV buffer. **(B)** The total photon flux (p/s) from EVs bound to the cells was quantified using IVS.

## Discussion

EV holds promising potential for diagnostics and therapeutic applications, yet the field lacks standardized fundamental technology, including but not exclusive to isolation methods, characterization, engineering, and functional analyses. The storage of EVs is one such technology critically affects their integrity, cargo molecule, and functions, influencing the subsequent therapeutic efficacy for clinical applications.^28,29^ In this study, we evaluated the effect of short-term storage using buffers containing BSA^19^ and trehalose^20^ based on previous studies, which complemented the results of PBS-HAT buffer reported by Görgens et al., who used HSA instead of BSA. Our study demonstrated that EV storage in PBS severely affects the cell culture media-derived bioactive EV integrity, cargo protection, and targeting functions following the short-term storage at -80 °C. In addition, the formation of larger particles by particle fusion or aggregation starts immediately after re-suspension in PBS. We demonstrated that our EV buffer prevented the significant loss of EVs and the formation of larger particles and maintained pDNA packaging capacity after 7-day storage at 4°C. Notably, engineered EVs displaying targeting molecules have preserved their function in vitro when stored in the EV buffer at 4 °C for seven days, where the EV buffer did not alter in vivo target capacity.

There is an urgent need for robust and standardized methodologies because the vast number of EV-related research papers published vary widely in source samples, experimental parameters, procedures, and settings, often not reproducible.^30^ Therefore, since the first publication of the minimal information for studies of extracellular vesicles (MISEV), the International Society for Extracellular Vesicles (ISEV) has been making continuous community efforts to recommend reporting requirements.^12,13^ For example, the initial condition widely adopted by the community was the isolated EV storage in PBS buffer at -80°C^15^, which has been extensively used, however, without supporting experimental evidence. The recent report comprehensively analyzing the effect of PBS preservations of small EVs at 4 °C, −20 °C, and −80 °C demonstrated that regardless of the temperature or duration, these conditions extensively damaged EVs, thus suboptimal.^31^ Several studies examined the influence of storage conditions on EVs, including temperature, freeze-thawing, lyophilization^32-34^, and various buffer conditions.^16,20,21^ The use of cryoprotectants, one of the commonly used storage methods for unstable biomaterials, notably improved stored EV quality.^16,20,21^ For example, Bosch *et al*. demonstrated that adding trehalose to PBS prevented aggregation and improved the size distribution and numbers with improved preservation of biological activity.^20^ In addition to the direct impact of buffers, EV adsorption to tube walls reduces particle recovery.^18^ Pre-coating storage tubes can reduce the loss due to this adsorption.^18^ The use of excipients, such as bovine serum albumin (BSA) and Tween 20, further improved EV preservation without the loss of wound healing function.^35^ Our data showed reduced particle counts and pDNA recovery (Fig 2A and 3C) from PBS-EVs while pDNA per particle was unchanged between PBS and EV buffer (Fig 3B), supporting the previous report of PBS-EV adsorption to tube walls.^35^ Note that our data revealed that the adsorption to tube walls occurs immediately after re-suspension in PBS to the fresh samples and, thus, should be avoided.

Particle size analysis is the most common and relatively simple analysis method to measure particle size and numbers, but current technology has limitations. The widely used particle counters, such as nanoparticle tracking analysis (NTA), a widely used method to assess particle counts and size distributions with mean or median peak values, overlook increases of large particles in fewer numbers and smaller size particles under the detection limit. Our data that extracted values of larger particles (>200 nm) presented evidence of particle fusion or aggregation within the PBS buffer with statistical means in both fresh and preserved samples (Fig 2B). Yet, these measures lack validation of individual EVs that cannot exclude the possibility of counting non-EV particles, such as transfection complex^36^, protein aggregates, or serum byproducts. Nevertheless, these measures from the current study could serve as one indicator of size variation among easily adaptable storage conditions.

Evaluation of engineered EVs should encompass the essential features for therapeutic applications in addition to physical properties, such as specificity and protection of exogenously introduced cargo after storage. Our data suggested the -80 °C storage in PBS or media lost the protective function from nucleases which is consistent with previous reports.^16,18,34^ Most of these reports have evaluated the protective functions of EVs by assessing the bioactivity of cargo molecules^37^, which is an indirect assessment and cannot rule out the possibility that the molecular complex without EV protection exerts functional activities. To our knowledge, this is the first report evaluating the packaging capacity of eEVs and their changes after storage in PBS or EV buffer by quantitatively assessing the DNA copy numbers before and after the nuclease treatment. In addition, we demonstrated that our EV buffers retain eEV protective function and targeting after storage at 4 °C or -80 °C, by in vitro binding assay (Fig 3A-C) and in vivo biodistribution assay (Fig 5 A and B). It is worth noting that the stronger bioluminescence signals from the binding assay are likely due to the impact of BSA supplementation and not the higher EV numbers. Previous reports demonstrated the effect of BSA on improved bioluminescence signals.^38^ Therefore, our result indicates that the binding capacity is not improved but not disturbed by BSA supplementation.

The limitation of this study is the short-term storage period of up to one week. The data presented here resulted from multiple data points highly reproducible from one experimental condition. Thus, it is critical to evaluate long-term storage using conditions with each experimental variable, such as isolation methods, EV engineering methods, and different source cells. Nevertheless, our particle size parameters obtained from commonly used NTA measurements are widely applicable and potentially serve as one indicator of eEV distortion following storage.

In conclusion, our study provides a comprehensive analysis of the short-term storage effects on media-derived eEV in PBS and EV storage buffer on EV size, particle numbers, protective function, and target activity, suggesting an optimal condition for short-term eEV storage. This report presents; (1) an eEV size distribution assessment using an overlooked NTA parameter, which led to the finding of the immediate deteriorating effects of PBS storage and freezing on eEVs; (2) the improved storage of our EV buffer; and (3) the method of EV protection assay using quantitative measures of their cargo DNA by qPCR. We believe our work will add evidence and value to standardizing conditions for EV storage, including buffers and temperatures and their analytical methods to generate reproducible research data toward progressing in the field for clinical translation, in addition to the previous efforts for improving EV storage conditions.

## Acknowledgement

We thank Dr. Alicia Withrow and the MSU Center for Advanced Microscopy for technical assistance in TEM. This work was partly supported by the startup fund for Dr. Harada provided by **Michigan State University** and a grant from the **Elsa U. Pardee Foundation**.

## Supplemental Figures

**Figure S1.**
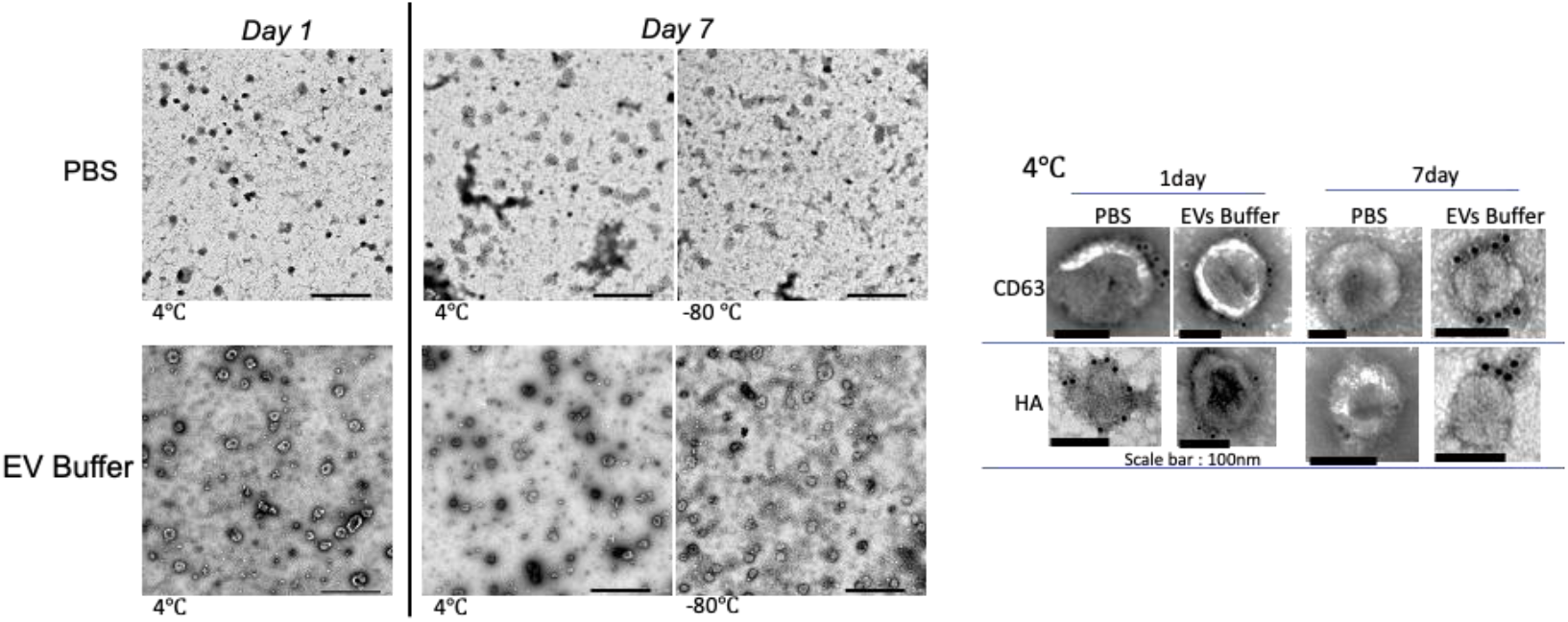
Transmission electron microscopy images of eEVs after the storages in PBS and the EV buffer showing gold-labeled HA and CD63 surface markers.

**Figure S2.**
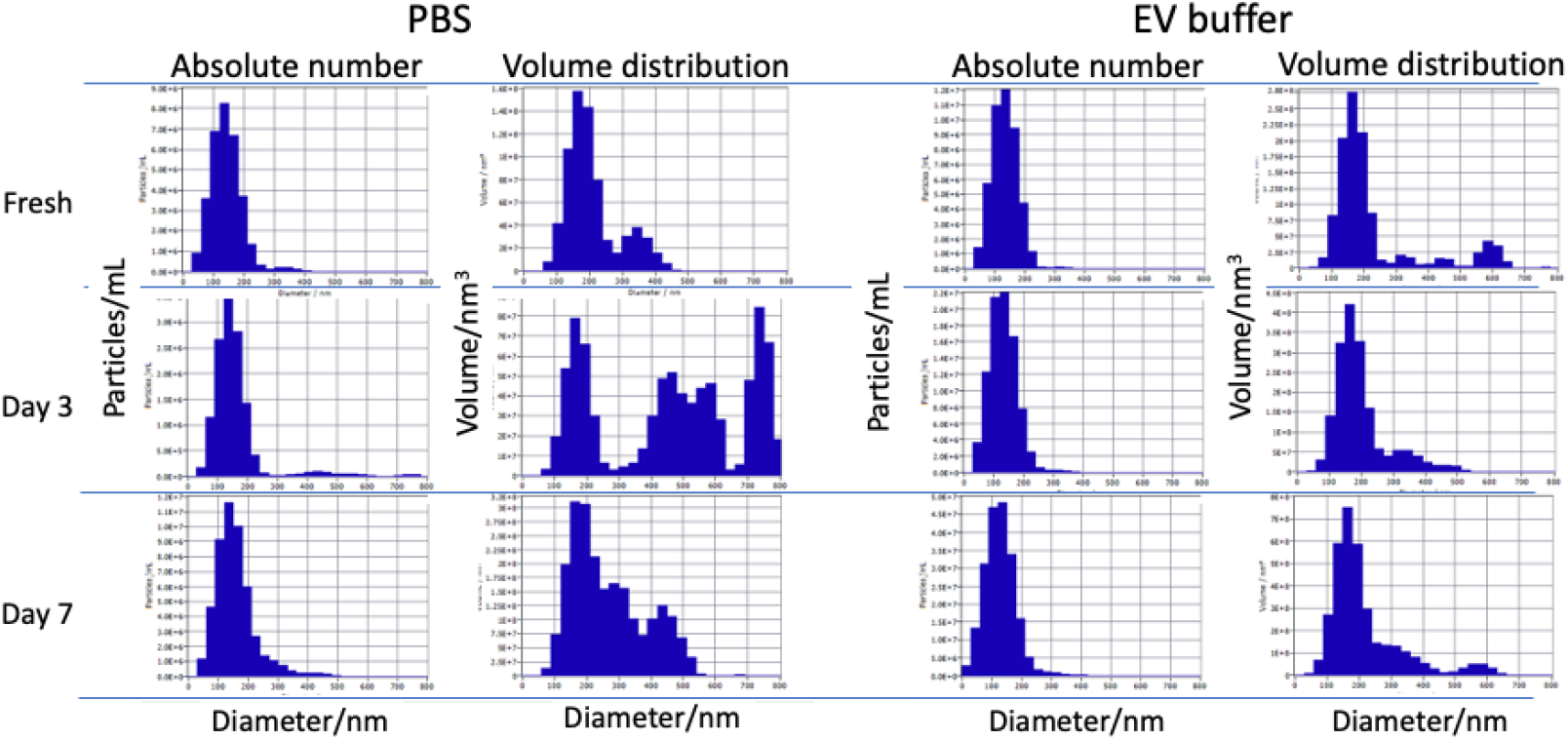
Typical examples of size distribution by different storage conditions. No significant change was observed on absolute number of EVs, PBS showed volume increase of large size particles than EV storage buffer.

**Figure S3.**
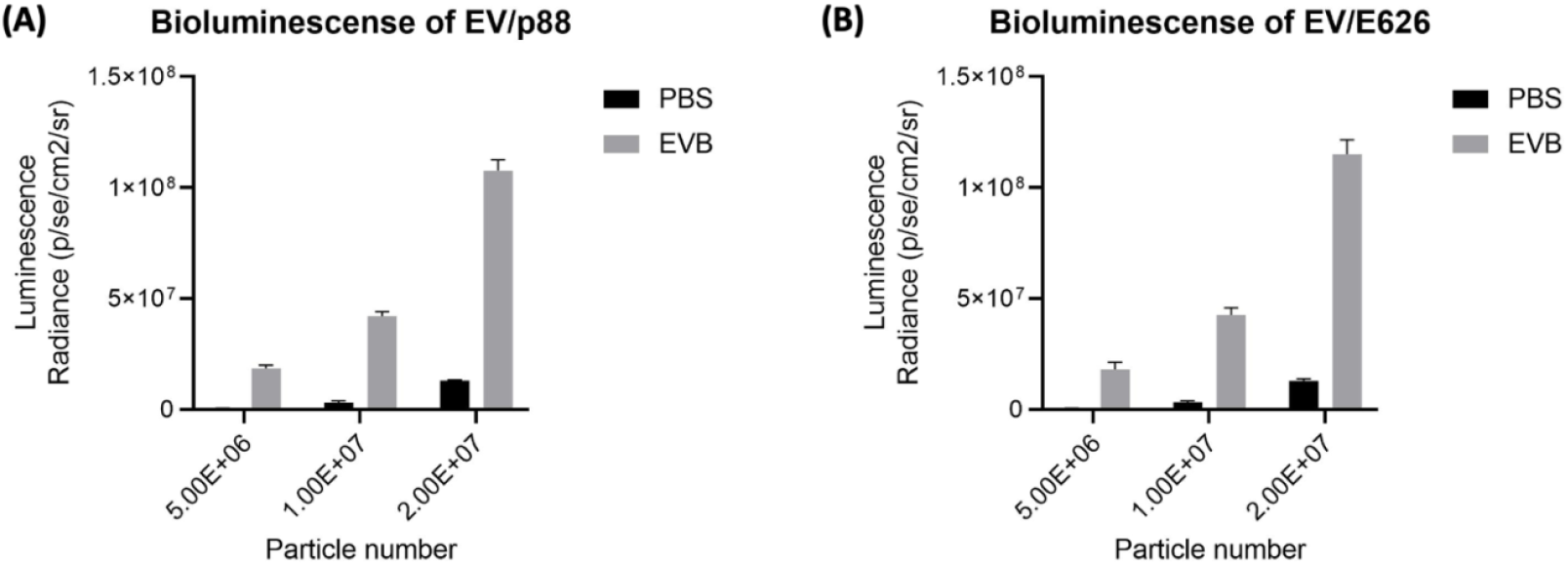
Comparison of eEV bioluminescence in PBS or EV buffer. Bioluminescence of 5×10^6^, 10^7^, and 2×10^7^ eEVs co-labeled with gLuc and p88 peptide **(A)** or E626 monobody **(B)** either in PBS or EV buffer were measured following the substrate addition. The total photon flux (p/s) from EVs bound to the cells by IVIS. The value represents the means ± SD (n = 3) in the graph.

